# Sapling: Accelerating Suffix Array Queries with Learned Data Models

**DOI:** 10.1101/2020.01.29.925768

**Authors:** Melanie Kirsche, Arun Das, Michael C. Schatz

## Abstract

**Motivation:** As genomic data becomes more abundant, efficient algorithms and data structures for sequence alignment become increasingly important. The suffix array is a widely used data structure to accelerate alignment, but the binary search algorithm used to query it requires widespread memory accesses, causing a large number of cache misses on large datasets.

**Results:** Here we present Sapling, an algorithm for sequence alignment which uses a learned data model to augment the suffix array and enable faster queries. We investigate different types of data models, providing an analysis of different neural network models as well as providing an open-source aligner with a compact, practical piecewise linear model. We show that Sapling outperforms both an optimized binary search approach and multiple existing read aligners on a wide collection of genomes, including human, bacteria, and plants, speeding up the algorithm by more than a factor of two while adding less than 1% to the suffix array’s memory footprint.

**Availability and implementation:** The source code and tutorial are available open-source at https://github.com/mkirsche/sapling.

**Supplementary Information:** Supplementary notes and figures are available online.

## 1. Introduction

Aligning sequencing reads to a reference genome or collection of genomes is a key component of many genomic analysis pipelines, including variant calling (Nielsen *et al.*, 2011), quantifying gene expression levels (RNA-seq) (Wang *et al.*, 2009), identifying DNA-protein binding sites (ChIP-seq) (Park, 2009) and several others (Soon *et al.*, 2013). Many techniques have been proposed to solve the read alignment problem in ways that are computationally efficient and robust to sequencing errors and true biological differences. Since finding inexact alignments is generally much slower than finding exact matches, a common approach is to use the seed-and-extend heuristic (Baeza-Yates and Perleberg, 1996). When using this heuristic, small segments of the read are used as seeds, and exact matches of these seeds are found using an algorithm for exact string matching. Then, the exact matches are used as candidate alignment sites, and each is scored based on how well the whole read aligns in the surrounding region. This heuristic has been shown to perform well in many genomic applications, and is used by a large number of leading short and long reads aligners including Star (Dobin *et al.*, 2013), Bowtie2 (Langmead and Salzberg, 2012), BWA-MEM (Li, 2013), NGMLR (Sedlazeck *et al.*, 2018) and many others. It is also used as a core routine for whole genome alignment (Marçais *et al.*, 2018) and many other applications (Altschul *et al.*, 1990).

The seed-and-extend heuristic relies on being able to quickly search for exact matches of seed sequences in the reference genome. The problem of finding these matches, called the exact substring search problem, has applications both within and outside of genomics (Charras and Lecroq, 2004). A number of data structures have been proposed to solve this problem by indexing the reference genome in such a way that the exact substring search problem can be solved quickly. These include suffix arrays (Manber and Myers, 1993), suffix trees (Weiner, 1973), hash tables (Karp and Rabin, 1987), and FM-indexes (Ferragina and Manzini, 2000). For genomic applications, suffix arrays are one of the key data structures for seed-and-extend algorithms used by Star (Dobin *et al.*, 2013), BLASR (Chaisson and Tesler, 2012), MUMMER4 (Marçais *et al.*, 2018) and others. The suffix array consists of the lexicographically ordered list of suffixes present in a string, and once constructed, a binary-search like algorithm can be used to quickly locate exact matches of query strings (Manber and Myers, 1993).

Learned index structures (Kraska *et al.*, 2017) are a technique for accelerating queries on a variety of data structures by leveraging patterns present in the particular dataset being processed. While classical data structures are asymptotically optimal, these runtime bounds are based on a worst-case analysis where it is assumed that the dataset has no specific patterns that can be exploited. However, many real-world datasets have learnable patterns, and learned index structures have been used in many different applications such as B-trees and Hash-maps (Kraska *et al.*, 2017). Additionally, learned index structures have previously been considered for read alignment using a modified FM-index (Ho *et al.*, 2019), although the source or implementation are not available and it was applied to a single dataset.

Here we present Sapling, an open-source algorithm which leverages learned index structures for accelerated read mapping. At its core, it uses suffix arrays, which we augment with a model of the particular genome that is being indexed. We evaluate two different types of data models - a neural network trained on the suffix array, as well as a compact piecewise linear model. We find that by using a data model, the core suffix array query time is reduced by more than a factor of two while only increasing the size of the data structure by less than 1% across a variety of genome sequences. We offer Sapling as both an open-source library for exact substring search and a standalone read aligner at https://github.com/mkirsche/sapling.

## 2. Methods

### 2.1 Suffix Array Search

For a text T of length n, let T[i] be the character in the ith position of T, and define a substring of T, T[i‥j], where 0 ≤ i ≤ j < n, as a string of characters T[i], T[i+1], …, T[j]. We define the exact substring search problem as follows: Given a text T of length n and a pattern P of length m, report all positions x in T such that T[x‥(x+m−1)] is equal to P. A naive algorithm of trying all possible values for P would take O(n * m) operations, which is infeasible for large texts, especially when many queries each need to be evaluated. In genomic applications where the text is a reference genome and the pattern is a genomic read a few properties generally hold: 1) The text is much (multiple orders of magnitude) larger than each query, and 2) The same text is used across multiple queries (typically many millions to billions of sequencing reads for a single genome). In an attempt to exploit these properties, several algorithms have been proposed which index the text on its own before any of the queries are considered, and then this index is used to reduce the number of possible alignment positions for every query.

One popular index is the suffix array. A suffix of T is defined as any substring T[i‥j] such that j = n − 1; that is, any substring which ends after the last character of T. Suffixes are related to substring search queries because any occurrence of a length-m pattern P at some position x in T corresponds to a suffix of T, T[x‥n−1], whose first m characters are exactly the string P. When the suffixes are considered in lexicographical order, all such suffixes starting with P will occur contiguously. This property of suffixes serves as the intuition behind the use of suffix arrays for exact substring search queries.

The suffix array is defined as an array of positions corresponding to the lexicographical order of suffixes in a given text. For a text T with n characters, we define the suffix array of T, SA_T_ to be a permutation of {0, ‥, n−1} such that SA_T_[i] is the start position in T of the ith suffix of T when the suffixes are sorted lexicographically. For example, in the text T = “CAT”, the sorted order of suffixes is {“AT”, “CAT”, “T”}, so SA_T_ = {1, 0, 2}. For any pattern P, each occurrence of P in T will be the prefix of some suffix of T, and since each such suffix starts with the characters in P, the start positions of the instances of P in T will occur consecutively in SA_T_. This reduces the problem of exact substring search to that of finding the range of suffix array positions [i, j] such that T[SA_T_[k]‥(SA_T_[k]+m−1)] = P for all integers k in [i, j]. These positions can be found using a binary search algorithm, which starts with an initial search space of [0, n−1] and repeatedly bisects the search space, querying the middle suffix to decide whether the suffixes starting with the characters in P occur in the first or second half, and recursively searching the half-sized space. The naive binary search algorithm, for a pattern of length m, requires O(log(n) * m) operations since each query requires a string comparison of up to m characters. However, a more efficient binary search algorithm specialized for the suffix array has been proposed which requires O(log(n) + m) operations. This exploits an auxiliary data structure called the longest common prefix array (LCP array) that stores the number of shared characters between the prefixes of consecutive suffixes (Manber and Myers, 1993).

### 2.2 A learned index structure for suffix arrays

When performing the binary search algorithm, each iteration requires checking the middle of the current search space. For large genomes, this means that consecutive iterations at the start of the algorithm correspond to distant array positions. Consequently, the algorithm has poor spatial locality and results in many cache misses. While the number of iterations is relatively small (~32 for a mammalian-sized genome), most of the memory accesses result in cache misses that are many times slower than memory accesses with cache hits - e.g., approximately 4ns to access from L1 cache vs 100ns to access from main memory on a modern Intel CPU (Levinthal, 2009). Therefore, we propose a method which uses a data model so that with a single memory lookup into the model and a small number of efficient arithmetic operations, the initial search space for binary search is significantly reduced, and the cache misses which occur at the beginning of the binary search algorithm can be mostly circumvented.

As described above, learned index structures have been used to replace or augment data structures with a data model which models some properties of the particular data being stored. In the case of suffix arrays, we define for a suffix array SA_T_ a true mapping T(x) which maps a k-mer x to the set of positions of the suffix array that correspond to suffixes starting with x. From the data, we learn a function P(x), a low-memory and arithmetically efficient approximation of T. Then, for a query k-mer Q, P(Q) gives an approximate position of where in the suffix array Q occurs. By performing this query on every k-mer in T, we can obtain a global error bound E on the predictions, which has the property that for any suffix in T, P(x) gives a position which is no more than E positions away from the nearest value in T(x). For a given k-mer x, we can compute P(x), and if x is present in the suffix array, there will be some suffix array position y in [P(x) − E, P(x) + E] such that the suffix starting at position SA_T_[y] starts with x, and this value of y can be computed using a binary search with an initial range of length 2E + 1 instead of length n (**Figure 1**). Therefore, we seek a model with three properties: the ability to perform predictions quickly, a low memory footprint, and a small error bound across genomes.

**Figure 1.**
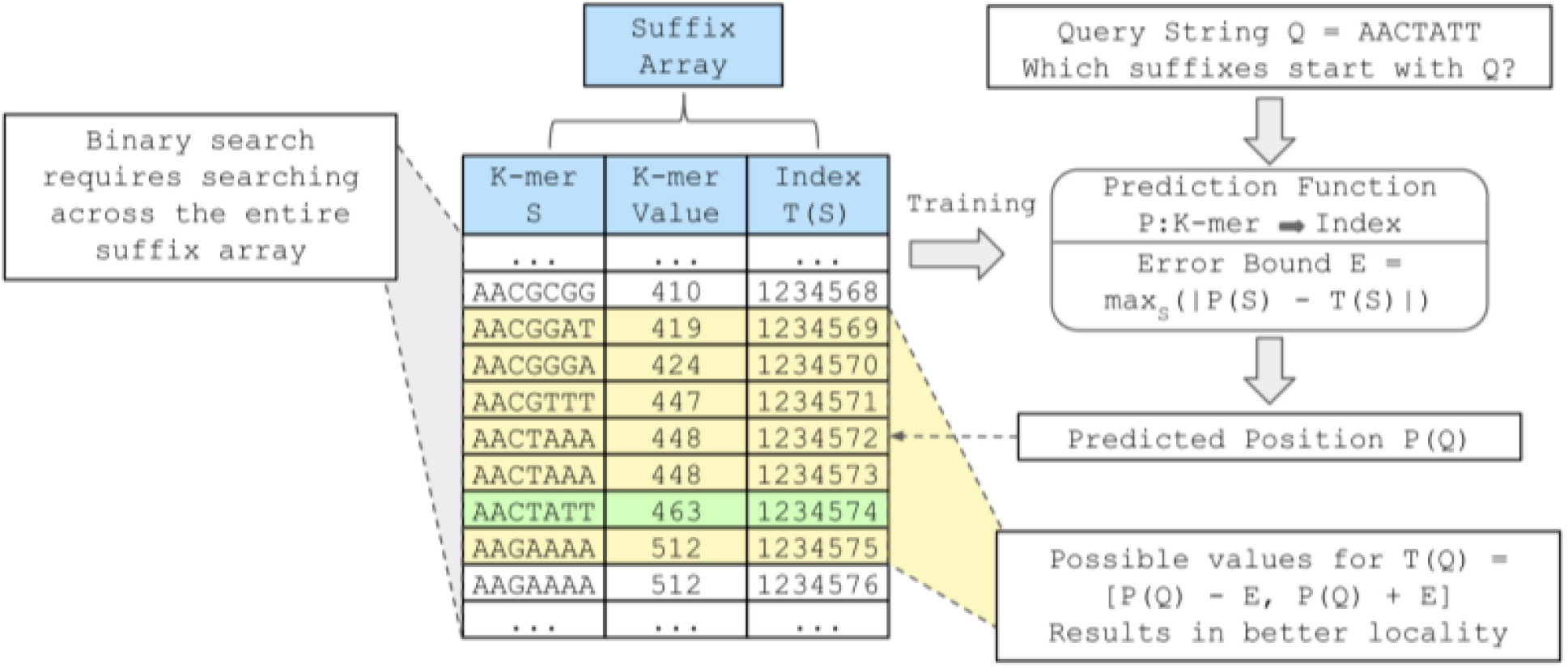
The suffix array lookup can be considered a prediction problem by defining a mapping T(S) which maps a k-mer S encoded as an integer K-mer value to each of the positions in the suffix array corresponding to suffixes starting with S. Learned index structures can approximate this mapping with a function P(S) mapping the k-mer value of each k-mer S to an estimated index, which is trained on the suffix array for a particular dataset using the (S, T(S)) pairs. The maximum error E across all k-mers in the string is computed so that when a particular k-mer Q is queried, if it is present in the string, then at least one of its suffix array positions falls in the range [P(Q) − E, P(Q) + E]. This smaller range can be used for the binary search lookup, resulting in better spatial locality.

### 2.3 Modeling with Artificial Neural Networks (ANNs)

The first method we explored for modeling the suffix array distribution was using an Artificial Neural Network (ANN) (Cybenko, 1989) to learn the true mapping T(x). In this approach, we trained ANNs on (k-mer value, suffix array position) pairs, with the goal of using the trained network to predict the approximate suffix array position of a given k-mer **(Figure 2a)**. To ensure that the function being learned is over numeric values, Sapling encodes each k-mer as its k-mer value - an integer with 2k bits. In this conversion, two bits are allocated for each of the k characters, with the two highest-order bits corresponding to the first character and the two lowest-order bits corresponding to the last character. The two bits for a given character are 00 if the character is “A”, 01 for “C”, 10 for “G”, and 11 for “T”. This encoding scheme ensures that any k-mer which comes lexicographically before another will have a smaller integer value, resulting in a simple, monotonically non-decreasing mapping.

**Figure 2.**
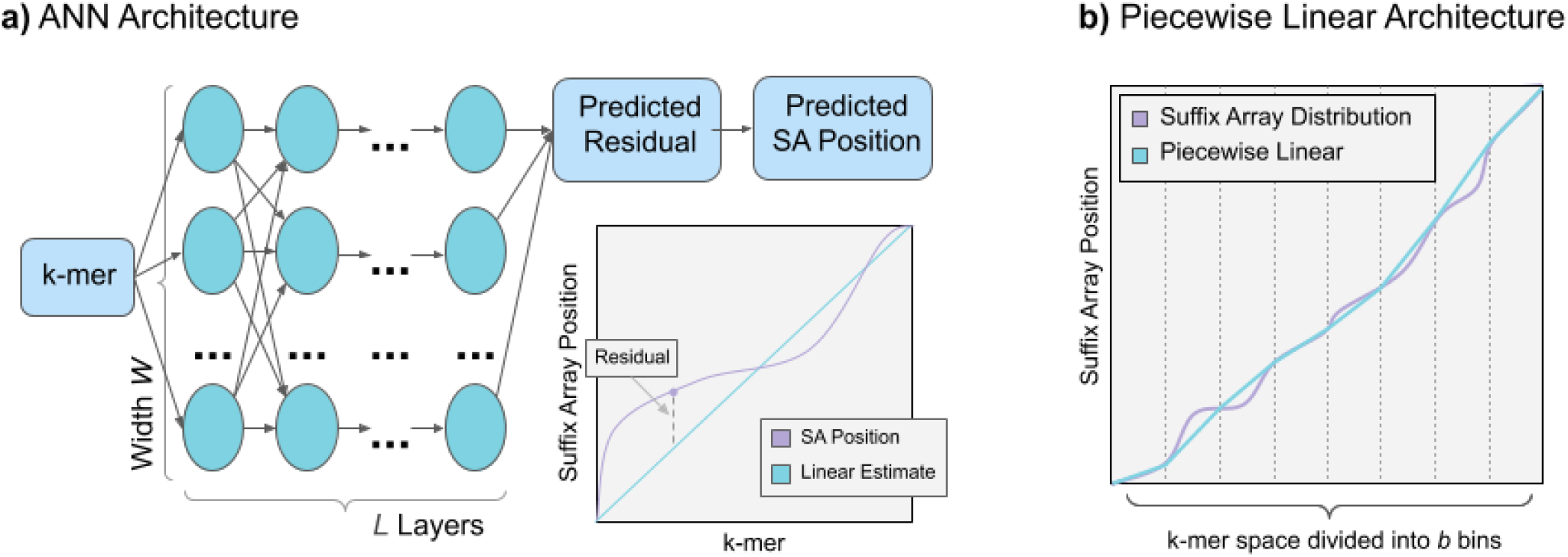
**a) Schematic diagram of ANN architecture.** An input k-mer encoded using a simple binary encoding scheme is passed to a fully connected ANN with L layers, each with width W. The output value from the ANN is the predicted residual value, which is then projected to the actual suffix array position using a linear transformation. In practice, we use multiple ANNs that each learn the distribution of a portion of the k-mer space (not shown). **b) Schematic diagram of Piecewise Linear model.** The piecewise linear model divides the space of possible inputs (k-mers encoded as integers) into b equal-sized intervals. It stores representative data points from each interval (those with the lowest k-mer values) and connects points in consecutive intervals with line segments. Then, when estimating the suffix array position for a particular k-mer, the linear function between that k-mer’s bin and the following bin is used to estimate the suffix array position.

For modeling, we first transform the suffix array positions into “residual values” - this detrending is performed by considering a straight line from the first k-mer to the last k-mer (i.e. fitting a linear function to the entire genome, such as plotted in **Figure 2a**), and then computing how each suffix array position differs from this line. The residual values are more easily learned by the ANN since the function will have a smaller range of values to consider. The input data is then unit scaled so that both the k-mers and the suffix array positions are within [0, 1]. We divide the input data into B equal-sized intervals, and an individual ANN is trained on each of them. For these neural nets, we used a basic “rectangular” architecture consisting of L layers, each with N nodes (aside from the single input node in the first layer and the single output node in the last layer). The networks were fully connected (each node in layer i passed input into every node in layer i+1), with no drop out, with a ReLU activation function (Ramachandran *et al.*, 2017) applied between layers.

The loss function used was mean squared error (the average of the square of the differences between the predicted suffix array residual position and the true value). We trained the model to minimize this loss function using the Adam optimizer with default PyTorch (Paszke *et al.*, 2019) hyper-parameters (learning rate 0.001, betas = [0.9, 0.999] and epsilon = 1e-8). The training for these models proceeds in epochs, during which the model’s ability to predict the input data is assessed and improved. During each epoch, the current model (using the parameters it has learned up to that point) makes predictions on the input data, and the mean squared error is computed. Based on this error, the parameters in the model are updated through a process called back propagation. To speed up training, we used a batch size of 64 - this means that the model makes predictions for 64 input values, the mean square error is calculated across these 64 predictions, and the model’s parameters are updated accordingly, before the next batch is loaded. The input data is shuffled at the start, so the batches do not contain consecutive data points.

For training, we set the maximum number of training epochs to be 200. All models were trained for at least 10 epochs, and after this initial period, if a reduction of 10% or more in the value of the loss function was not achieved during the last 10 epochs, the training procedure was terminated to limit wasted work. When the training for a particular neural network ended, the best model across all training epochs was kept and used to predict the suffix array positions for all k-mers in the network’s corresponding interval of k-mer values.

### 2.4 Modeling with Piecewise Linear Functions (PWL)

An alternative data model we explored is a piecewise (PWL) linear model. In this model, the space of all 4^k^ possible k-mers is subdivided into a fixed number b equally-sized intervals, where b is a power of 2 to allow fast calculation of which interval each k-mer falls into **(Figure 2b)**. Then, for each interval, the lexicographically earliest k-mer from the genome which is present in that interval is stored along with its corresponding suffix array position. While this idea of “marker” k-mers to limit the range of the suffix array to search has been used previously (Dobin *et al.*, 2013), Sapling improves upon this approach by interpolating the exact suffix array position of the entire k-mer, giving an even smaller interval of candidate positions. In the algorithm used by Sapling, the prediction P(s) is computed as follows:

1. Calculate which interval x is in by shifting its value right by b bits.
2. Look up the pair (x_1_, y_1_) corresponding to the earliest k-mer in the same interval and the pair (x_2_, y_2_) corresponding to the earliest k-mer in the next interval.
3. Construct a line segment between (x_1_, y_1_) and (x_2_, y_2_), and output the y-value which corresponds to an x-value of s.

This simple model allows very efficient queries consisting of looking up two pairs which are adjacent in memory followed by a small number of arithmetic operations. The memory footprint is parameterized on the number of intervals, storing two 64-bit integers per interval, and we show that even with a relatively small number of intervals, small error bounds can be achieved across different genomes. For these reasons we use this data model in our implementation.

### 2.5 PWL Implementation

When dividing the space of possible k-mers into buckets, the partitioning is done in such a way that each group has the same number of possible k-mers. However, in practice, due to varying k-mer frequencies, it is possible for some buckets to have particularly small or large sections of the suffix array contained in them. The buckets with many points often have particularly poor predictions, and this causes the maximum errors to be much worse than the median errors or even the 95th percentile errors (see **Results**). To avoid binary searching over a range which is almost always much larger than necessary, Sapling uses an additional cutoff. Once the predictions have been made for every k-mer in the genome, in addition to storing the maximum error in each direction, Sapling also stores the 95th percentiles of the errors in each direction. Then, when searching for a particular k-mer given its predicted position, rather than immediately executing the binary search algorithm, Sapling first checks the position corresponding to an error equal to the 95th percentile in the appropriate direction. Then, in 95% of cases, the size of the search range can be immediately reduced to the 95th percentile error, which is much smaller than the maximum error, further improving performance.

When using Sapling, it is assumed that the size of k-mers used when constructing the index is equal to the length of the k-mers being queried. However, it is possible that for some applications, the index will be searched for queries of varying lengths. The suffix array prediction function can be evaluated for such strings without rebuilding the model as follows:

➢ If the query length q is less than the Sapling k-mer size k: Pad the end of the query with A’s (the k-mer value can be padded in this way quickly by bit-shifting the k-mer value 2*(k−q) bits to the left).
➢ If the query length q is greater than the Sapling k-mer size k: Let the k-mer value of the length-k prefix of the query be v. Then, set the k-mer value of the query as a floating-point value between v and v+1 based on the remaining characters and evaluate the piecewise linear function at that value.

Sapling is available as open-source software on Github, and provides a succinct library for constructing the piecewise linear data model and using it to perform suffix array lookups. We also implemented a simple seed-and-extend aligner as a proof-of-concept which uses Sapling for seeding and the Striped-Smith-Waterman algorithm (Zhao *et al.*, 2013) for extending seeds into full alignments. This aligner accepts fasta and fastq formatted files as input and outputs alignments in SAM format (Li *et al.*, 2009).

## 3. Results

### 3.1 Suffix Array Distribution

We tested Sapling on six diverse reference genome sequences: *E. coli*, *C. elegans*, *S. lycopersicum* (tomato), human (both chromosome 1 in isolation and the full human reference), and *T. aestivum* (wheat) (**Supplemental Table 1, Supplemental Figure 1)**. While the function we are trying to approximate is monotonically non-decreasing, there are many potential functions that can emerge based on the composition of the suffix array. While the suffix array for a random string will result in approximately a straight line, repetitiveness and biological selection against certain sequences (Herold *et al.*, 2008) can drastically affect the nature of the function (**Supplementary Figures 2-3**). Therefore, we investigated the true suffix array position functions for each of these genomes to ensure that the functions are learnable across species. **Figure 3** shows the true Suffix Array Distributions for each of the six reference genomes listed above.

**Figure 3.**
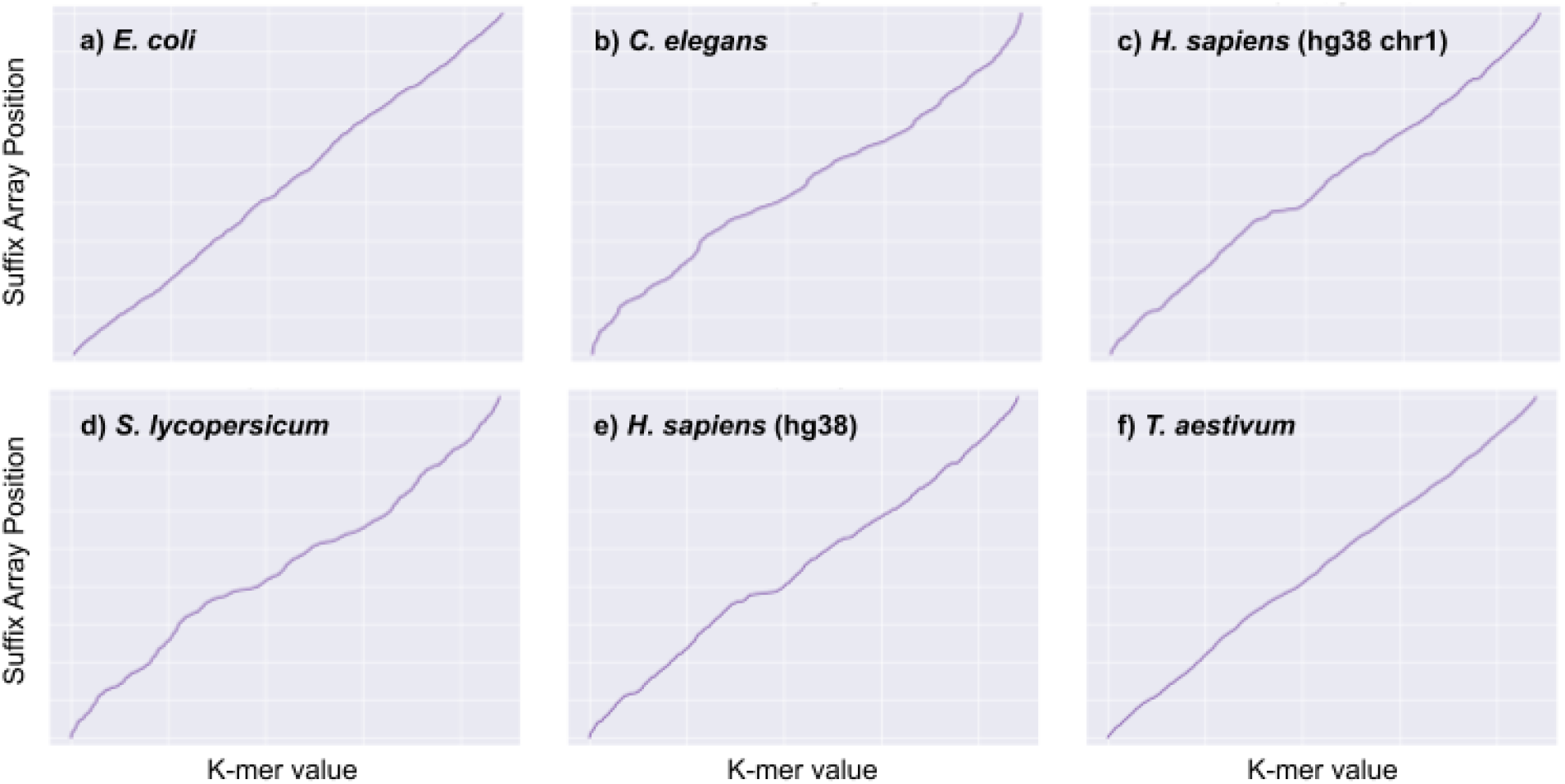
Suffix Array Distribution for 6 genome sequences: *E. coli*, *C. elegans* (nematode), *H. sapiens* (chr1), *S. lycopersicum* (tomato), *H. sapiens* (all of hg38), and *T. aestivum* (wheat).

### 3.2 Model Training and Accuracy

In testing the feasibility of different models, we measured the prediction accuracy of several potential PWL and ANN models on human chromosome 1 (**Supplemental Tables 2 & 3)**. **Table 1** describes the characteristics of a selection of model architectures as well as their memory footprints. For the ANN, most bins are trained within 40-60 epochs, although a few particularly complex bins require up to 180 epochs until convergence (**Supplemental Figure 4**). For each model, we calculated the prediction error for every k-mer present in the genome, defined as the absolute difference between the predicted suffix array position and the nearest position which corresponds to a suffix starting with the query. The mean, median, and maximum errors were computed both within each bin and genome-wide. By studying each bin individually, we were able to highlight cases where the learned function modelled the suffix array position function particularly well or poorly (**Supplementary Figures 5 & 6**). In particular, for all of the genomes we studied, the first and last bins had particularly high prediction errors caused by the high relative frequencies of homopolymer A and T sequences in the genomes that challenged the PWL model. We found that increasing the width of the ANN used for each bin in the model resulted in improved performance, without adding much overhead. However, we found that while increasing the depth (number of layers) of each ANN in the model resulted in performance increases, it added significant memory overhead. This leads us to conclude that utilizing shallower, wider nets is the most efficient way to approach this problem. Overall, the PWL model had improved median and 95% percentile accuracy compared the ANN model, especially when considering the memory overhead involved, although the ANN model had a lower maximum error.

**Table 1.** Summary of performance and model complexities for several PWL and ANN architectures.

| Model Type | Piecewise Linear | Piecewise Linear | Piecewise Linear | Neural Network | Neural Network | Neural Network |
| --- | --- | --- | --- | --- | --- | --- |
| Number of Buckets | 16k | 256k | 2m | 1k | 16k | 16K |
| Width x Depth | N/A | N/A | N/A | 32 x 1 | 32 x 1 | 128 x 2 |
| Median Error | 899 | 68 | 14 | 900 | 131 | 56 |
| 95th Percentile Error | 7,658 | 1,579 | 653 | 4238 | 853 | 463 |
| Maximum Error | 263,165 | 180,453 | 135,664 | 45,839 | 24,081 | 13,264 |
| Memory Overhead | 256 KB | 4 MB | 32 MB | 8 MB | 131 MB | 1245 MB |

### 3.3 Runtime analysis

Based on the above results, we implemented Sapling to use the PWL data model to accelerate suffix array queries. We then compared the performance of Sapling using different numbers of bins to a number of existing alignment algorithms (**Figure 4, Supplemental Tables 4 & 5, Supplemental Note 1**). For this, we implemented a string-optimized binary search, the asymptotically optimal algorithm for searching a suffix array (Manber and Myers, 1993). We also ran the widely-used Bowtie (Langmead *et al.*, 2009) and Mummer4 (Marçais *et al.*, 2018) short read aligners in their exact-matching modes (**Supplementary Note 1**) to obtain a fair comparison to Sapling’s performance. For each aligner, we measured the amount of time needed to perform 50 million queries on the human genome, where each query is a random 21-mer which is known to occur in the genome, ignoring the time required for indexing. All software was run on a single core of an Intel(R) Xeon(R) CPU E7-8860 server at 2.20 GHz with 1 TB of RAM. For the runtime experiments we used a tmpfs ramdisk to minimize the amount of IO-based latency. As expected, we see the runtime performance of Sapling improves as the number of bins increases. In an ideal case, with a perfect prediction function, the number of suffix arrays lookups would be decreased from log_2_(n) - approximately 32 for the human genome - to a single lookup at the predicted position. Our model is able to reduce the search range to a few thousand rows, reducing the number of lookups to about 10 for most queries. This results in an algorithm which is more than 3 fold faster than the string-optimized binary search and nearly 6.5 fold faster than bowtie when used with the largest number of bins.

**Figure 4.**
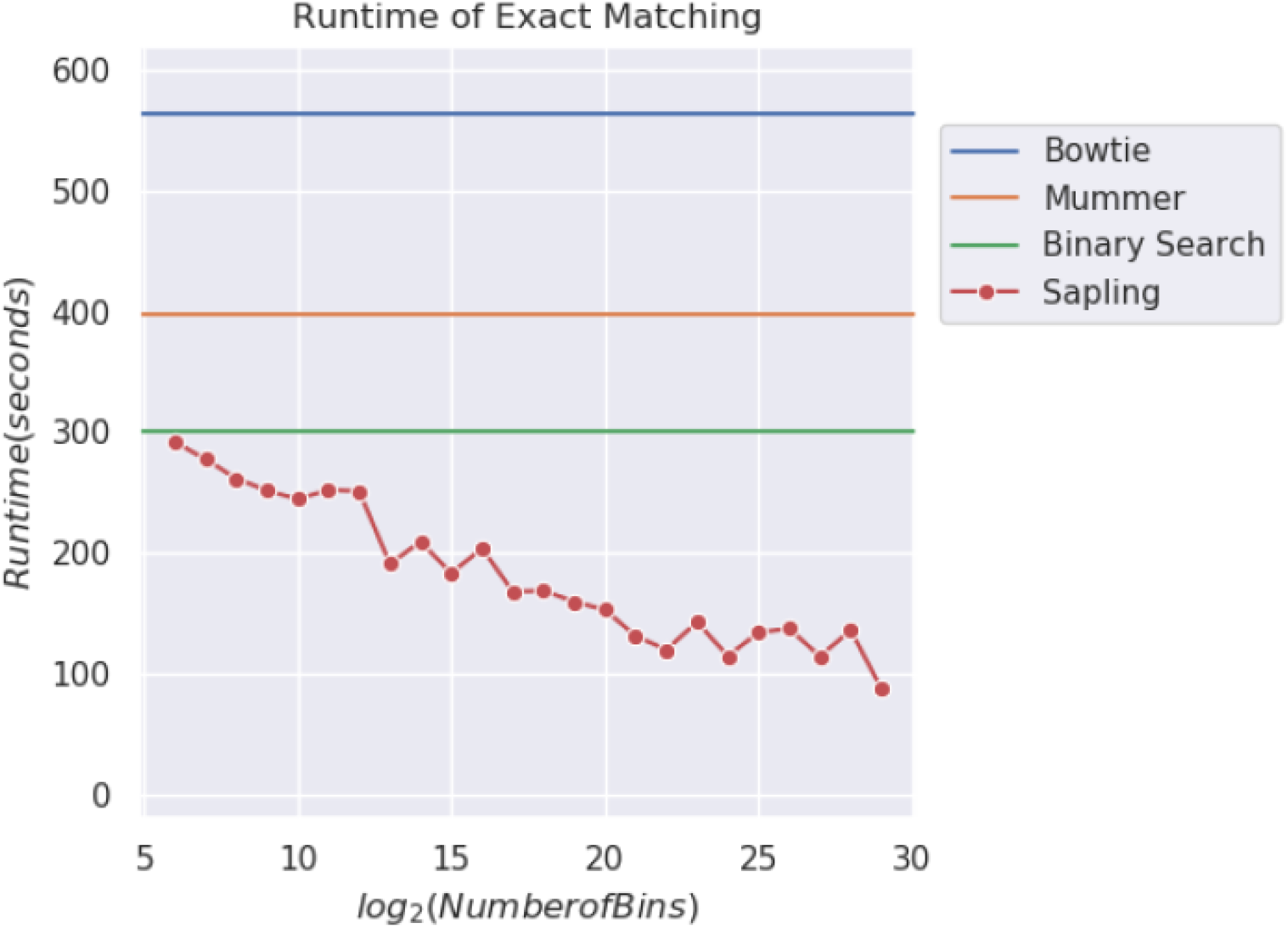
Runtime of different methods to locate 50 million 21-mers in the human genome. The queries were sampled randomly from those which occur at least once in the human genome, and the same queries were used for all methods. In addition to running Bowtie, Mummer4, and a string-optimized binary search algorithm, Sapling was run with several different settings, limiting the number of buckets (and therefore the memory overhead) to various proportions of the size of the human genome.

In addition to measuring across model architectures and between different aligners, we also measured how well the runtime of Sapling scales when the genome size is increased. To measure this, we ran Sapling on six different reference genomes of different sizes, and for each genome measured the amount of time required to query five million random k-mers which are present in the genome. We performed a similar experiment for the string-optimized binary search. We find that as the genome size increases, the reduction in runtime from using a data model increases substantially (**Figure 5, Supplemental Tables 6 & 7**).

**Figure 5.**
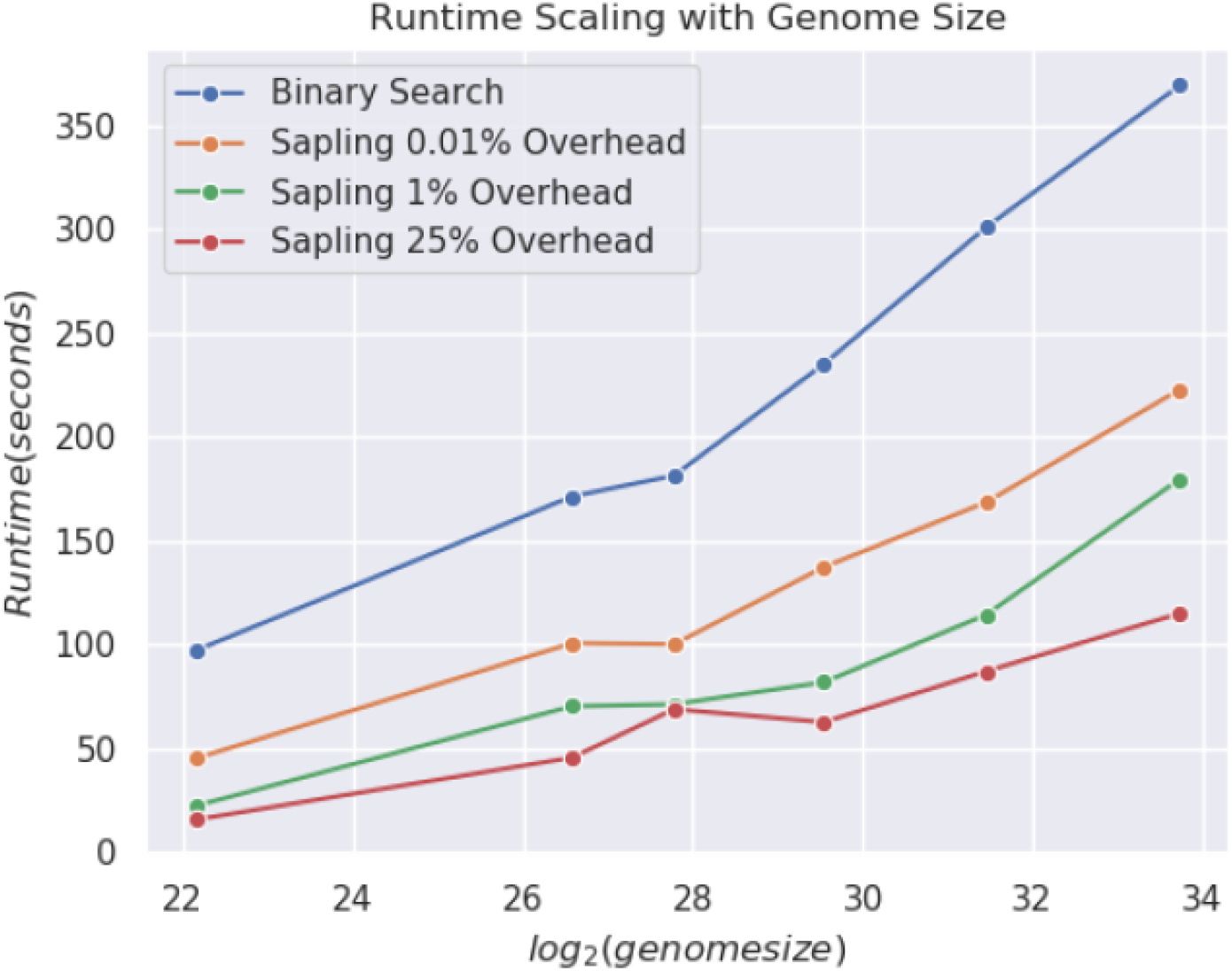
Runtime of Sapling and binary search across six different genomic sequences using 0.01%, 1% or 25% space overhead.

## 4. Discussion

In this paper, we presented Sapling, a novel algorithm for quickly performing suffix array lookups for use within read alignment and genome alignment algorithms. Sapling uses the idea of learned index structures to model the contents of the suffix array as a function rather than as a data structure, and uses a practical piecewise linear model to efficiently approximate this function. Using this method shows significant improvement in the runtime of querying many different genomes, demonstrating that even a simple low-memory piecewise linear approximation of the suffix array position function is sufficient for achieving several-fold improved performance compared to existing tools with modest space overhead. As read and genome alignment is performed on even larger genomes and larger collections of genomes, the need for efficient substring search algorithms becomes even more pressing, and Sapling will be able to scale better to large reference sizes than existing query algorithms.

While this work demonstrates the potential for learned index structures in a very important and widely used genomic application, there remain many possible avenues for future development. Presently, the prototype read aligner uses a basic seed-and-extend implementation that requires additional development to make it competitive with existing aligners for inexact alignment. In addition, Sapling could be used for modeling other full text index data structures, especially sparse versions of the suffix array (Vyverman *et al.*, 2013) or the FM-index, or other data structures entirely. Finally, read alignment is just one of the many data-intensive problems in genomics that requires the efficient use of large data structures. We are investigating other genomic applications of the learned index structures paradigm, including optimized graph representations for genome and pan-genome assembly, optimized variant databases, and other data intensive problems.

## Acknowledgements

We would like to thank Alex Dobin and Benjamin Langmead for their helpful discussions. This research project was conducted using computational resources at the Maryland Advanced Research Computing Center (MARCC).

## Funding

This work was supported by the National Science Foundation [DBI-1350041, IOS-1445025, IOS-1732253, and IOS-1758800 to MCS].

## Conflict of Interest

none declared.

