## Supplementary Material for "Sapling: Accelerating Suffix Array Queries with Learned Data Models"

### Supplemental Materials

**Supplemental Table 1. Genome sequences analyzed in this study.** 2

**Supplemental Table 2. Prediction errors for the piecewise linear data model.** 3

**Supplemental Table 3. Predictions errors for ANN modeling.** 4

**Supplemental Table 4. Runtime of different methods querying the human genome.** 5

**Supplemental Table 5. Runtime of Sapling on the human genome.** 6

**Supplemental Table 6. Runtime of binary search on six genomic sequences.** 7

**Supplemental Table 7. Runtime of Sapling on six genomic sequences.** 8

**Supplemental Figure 1. K-mer Uniqueness Ratios of Genomes.** 9

**Supplemental Figure 2. Suffix array distributions for simulated repetitive genomes.** 10

**Supplemental Figure 3. Suffix array distributions for simulated genomes with different nucleotide compositions.** 11

**Supplemental Figure 4. Distribution of the number of epochs until convergence.** 12

**Supplemental Figure 5. Examples of PWL models for selected bins in human chr1.** 13

**Supplemental Figure 6. Examples of ANN performance for selected bins in human chr1.** 14

**Supplemental Note 1. Commands used for running different aligners** 15

**Supplemental Table 1. Genome sequences analyzed in this study.**

Note that all ‘N’ characters were removed from the sequences prior to indexing.

| **Species Name** | **Genome Size** | **Accession** |
|---|---:|---:|
| *E. coli* K-12 substr. MG1655 | 4,641,652 | GCA_000005845.2 |
| *C. elegans* | 100,286,401 | GCA_000002985.3 |
| *S. lycopersicium* | 782,475,302 | SL4.0 (https://www.biorxiv.org/content/10.1101/767764v1) |
| *H. sapiens (hg38 chr1)* | 230,481,012 | GCA_000001405.15 |
| *H. sapiens (hg38 whole genome)* | 2,934,876,451 | GCA_000001405.15 |
| *T. aestivum* | 14,271,578,887 | GCA_900519105.1 |

**Supplemental Table 2. Prediction errors for the piecewise linear data model.**

This table shows a number of error statistics for different numbers of bins when using the piecewise linear model to predict the suffix array positions of all 21-mers in human chromosome 1.

| Number of Bins | Max Error | 95th Percentile | Median Error | Mean Error | Median of medians | Max of medians |
|---:|---:|---:|---:|---:|---:|---:|
| 2^0 | 12,987,798 | 11,299,286 | 6,899,176 | 6,502,090 | 6,899,176 | 6,899,176 |
| 2^10 | 426,608 | 59,211 | 11,590 | 19,578 | 9,429 | 273,216 |
| 2^14 | 263,165 | 7,658 | 899 | 2,533 | 537 | 163,013 |
| 2^18 | 180,453 | 1,579 | 68 | 627 | 27 | 126,588 |
| 2^20 | 150,267 | 828 | 19 | 395 | 7 | 108,158 |
| 2^21 | 135,664 | 653 | 14 | 335 | 4 | 103,207 |
| 2^22 | 124,792 | 486 | 6 | 270 | 2 | 88,840 |
| 2^23 | 111,458 | 392 | 4 | 234 | 1 | 84,460 |
| 2^24 | 102,668 | 314 | 2 | 194 | 1 | 71,826 |
| 2^25 | 90,667 | 257 | 2 | 170 | 1 | 67,470 |
| 2^26 | 83,653 | 221 | 1 | 142 | 0.5 | 56,666 |
| 2^27 | 72,889 | 187 | 1 | 127 | 0 | 52,476 |

**Supplemental Table 3. Predictions errors for ANN modeling.**

This table shows a number of error statistics for different ANN architectures to predict the suffix array positions of all 21-mers in human chromosome 1.

| # Buckets | # Nodes | # Layers | Median of Means | 95th %tile of Means | Max of Means | Median of Medians | 95th %tile of Medians | Max of Medians | Median of Maxes | 95th %tile of Maxes | Max of Maxes | Total Size (MB) |
|---|---|---|---|---|---|---|---|---|---|---|---|---|
| 2^10 | 8 | 1 | 3,082 | 9,259 | 27,850 | 2,432 | 8,425 | 26,627 | 10,709 | 26,532 | 91,606 | 8 |
| | 32 | 1 | 1,194 | 2,559 | 11,042 | 892 | 1,792 | 4,752 | 5,434 | 13,088 | 45,839 | 8 |
| | 128 | 1 | 713 | 1,879 | 10,165 | 520 | 1,102 | 4,065 | 3,775 | 12,551 | 50,428 | 9 |
| | 8 | 2 | 1,888 | 5,353 | 235,792 | 1,406 | 4,433 | 229,429 | 7,611 | 19,372 | 523,256 | 9 |
| | 32 | 2 | 683 | 1,674 | 9,174 | 492 | 1,082 | 3,977 | 3,799 | 12,120 | 38,627 | 13 |
| | 128 | 2 | 435 | 1,318 | 8,868 | 308 | 696 | 3,677 | 3,257 | 12,144 | 36,495 | 76 |
| | 8 | 4 | 1,197 | 4,904 | 106,727 | 835 | 4,190 | 102,676 | 5,730 | 21,838 | 250,781 | 10 |
| | 32 | 4 | 435 | 1,318 | 8,868 | 308 | 696 | 3,677 | 3,257 | 12,143 | 36,495 | 22 |
| | 128 | 4 | 332 | 1,336 | 9,790 | 208 | 622 | 4,976 | 3,217 | 12,332 | 51,049 | 204 |
| 2^14 | 8 | 1 | 302 | 1,228 | 3,494 | 245 | 1,116 | 3,516 | 967 | 3,394 | 34,965 | 131 |
| | 32 | 1 | 164 | 757 | 3,458 | 128 | 494 | 3,452 | 603 | 2,699 | 24,081 | 131 |
| | 128 | 1 | 105 | 641 | 3,463 | 29 | 334 | 3,452 | 432 | 2,475 | 22,944 | 147 |
| | 256 | 1 | 92 | 625 | 3,491 | 70 | 325 | 3,452 | 394 | 2,453 | 9,079 | 180 |
| | 512 | 1 | 86 | 630 | 3,817 | 66 | 370 | 3,813 | 373 | 2,439 | 23,573 | 229 |
| | 8 | 2 | 211 | 987 | 35,024 | 163 | 826 | 34,404 | 736 | 3,002 | 67,349 | 147 |
| | 32 | 2 | 113 | 586 | 3,456 | 85 | 324 | 3,453 | 467 | 2,341 | 8,687 | 212 |
| | 128 | 2 | 71 | 505 | 3,448 | 52 | 252 | 3,452 | 325 | 2,224 | 13,264 | 1,245 |
| | 8 | 4 | 154 | 1,015 | 43,595 | 115 | 808 | 47,065 | 597 | 3,111 | 83,259 | 163 |
| | 32 | 4 | 72 | 463 | 3,460 | 53 | 205 | 3,452 | 349 | 2,199 | 7,218 | 360 |
| | 128 | 4 | 70 | 390 | 3,453 | 30 | 183 | 3,452 | 221 | 1,937 | 7,061 | 3,342 |
| 2^18 | 128 | 1 | 128 | 129 | 223 | 2 | 18 | 220 | 61 | 127 | 642 | 2,359 |

**Supplemental Table 4. Runtime of different methods querying the human genome.**

Each method was used to locate 50 million 21-mers that are known to occur in the genome. Speed up is computed relative to the suffix array binary search. **Supplemental Table 5** displays additional results for Sapling.

| Tool | Runtime (s) | Speed up over binary search |
|---|---:|---:|
| Bowtie | 564 | .53x |
| Mummer4 | 396 | .76x |
| Suffix Array Binary Search | 301 | 1x |
| Sapling (0.01% memory overhead) | 169 | 1.78x |
| Sapling (1% memory overhead) | 114 | 2.64x |
| Sapling (25% memory overhead) | 87 | 3.46x |

**Supplemental Table 5. Runtime of Sapling on the human genome.**

Each experiment was repeated 3 times to minimize the impact of contention on the server. The minimum recorded value is highlighted in green. Each run evaluates the time required to query 50 million randomly selected 21-mers known to occur in the genome.

| $\log_2$ (number of bins) | Trial 1 | Trial 2 | Trial 3 |
|---|---|---|---|
| 6 | 305.372 | 291.782 | 294.436 |
| 7 | 317.987 | 277.642 | 316.418 |
| 8 | 261.552 | 348.806 | 288.447 |
| 9 | 251.606 | 263.528 | 253.346 |
| 10 | 350.099 | 244.442 | 335.311 |
| 11 | 254.17 | 252.177 | 259.045 |
| 12 | 266.156 | 309.023 | 251.159 |
| 13 | 190.957 | 205.078 | 194.274 |
| 14 | 209.15 | 245.473 | 229.7 |
| 15 | 241.021 | 183.053 | 183.528 |
| 16 | 203.781 | 262.426 | 218.109 |
| 17 | 208.467 | 167.81 | 204.386 |
| 18 | 169.03 | 168.62 | 170.254 |
| 19 | 159.528 | 170.518 | 166.87 |
| 20 | 187.687 | 152.803 | 184.484 |
| 21 | 199.157 | 131.041 | 183.282 |
| 22 | 189.45 | 119.709 | 180.804 |
| 23 | 142.572 | 147.85 | 222.873 |
| 24 | 175.033 | 114.502 | 142.399 |
| 25 | 181.534 | 133.928 | 185.748 |
| 26 | 143.922 | 141.017 | 137.158 |
| 27 | 118.229 | 114.152 | 136.461 |
| 28 | 162.134 | 135.96 | 142.399 |
| 29 | 126.938 | 127.629 | 87.2305 |

**Supplemental Table 6. Runtime of binary search on six genomic sequences.**

Each experiment was repeated 3 times to minimize the impact of contention on the server. The minimum recorded value is highlighted in green. **Supplemental Table 7** shows the results for these experiments with Sapling.

| Genome | Runtime Trial 1 (seconds) | Runtime Trial 2 (seconds) | Runtime Trial 3 (seconds) |
|---|---|---|---|
| *E.coli* | 98.3190 | 97.197 | 98.9074 |
| *C. elegans* | 171.466 | 173.316 | 171.586 |
| *H. sapiens* (hg38, chr1) | 184.135 | 182.082 | 181.537 |
| *S. lycopersicum* | 237.823 | 236.077 | 235.254 |
| *H. sapiens* (hg38) | 481.586 | 336.127 | 301.056 |
| *T. aestivum* | 369.465 | 384.347 | 372.748 |

**Supplemental Table 7. Runtime of Sapling on six genomic sequences.**

Each experiment was repeated 3 times to minimize the impact of contention on the server. The minimum recorded value is highlighted in green.

| Genome | Sapling Overhead | Runtime Trial 1 (seconds) | Runtime Trial 2 (seconds) | Runtime Trial 3 (seconds) |
|---|---|---|---|---|
| *E. coli* | 0.01% | 36.4275 | 42.4027 | 45.2589 |
| | 1% | 22.7368 | 22.4503 | 22.6046 |
| | 10% | 16.4947 | 16.9750 | 16.5921 |
| | 25% | 17.1012 | 16.3602 | 15.7859 |
| *C. elegans* | 0.01% | 100.803 | 100.803 | 100.809 |
| | 1% | 85.3959 | 90.761 | 70.4513 |
| | 10% | 72.4352 | 64.8205 | 62.9823 |
| | 25% | 62.7908 | 45.6998 | 62.5775 |
| *H. sapiens* (hg38, chr1) | 0.01% | 124.404 | 100.316 | 132.331 |
| | 1% | 71.4348 | 71.6753 | 71.3830 |
| | 10% | 56.3261 | 56.5146 | 55.3909 |
| | 25% | 69.0131 | 81.8918 | 69.7210 |
| *S. lycopersicum* | 0.01% | 137.476 | 181.851 | 151.527 |
| | 1% | 88.0505 | 82.1407 | 82.0073 |
| | 10% | 94.6657 | 99.0286 | 102.249 |
| | 25% | 62.7804 | 67.3399 | 93.0033 |
| *H. sapiens* (hg38) | 0.01% | 169.03 | 168.62 | 170.254 |
| | 1% | 175.033 | 114.502 | 142.399 |
| | 10% | 162.134 | 135.960 | 142.399 |
| | 25% | 126.938 | 127.629 | 87.2305 |
| *T. aestivum* | 0.01% | 222.989 | 273.322 | 232.971 |
| | 1% | 179.87 | 187.326 | 281.909 |
| | 10% | 123.117 | 132.654 | 123.919 |
| | 25% | 136.917 | 181.883 | 115.208 |

**Supplemental Figure 1. K-mer Uniqueness Ratios of Genomes.**

For each of the genomes we used, we measured their repetitiveness by plotting how their k-mer uniqueness ratio (proportion of k-mers on the forward strand which occur exactly once) changes as a function of the k-mer length k. If the proportion of unique k-mers grows slowly as k increases, this indicates the presence of long repeats.

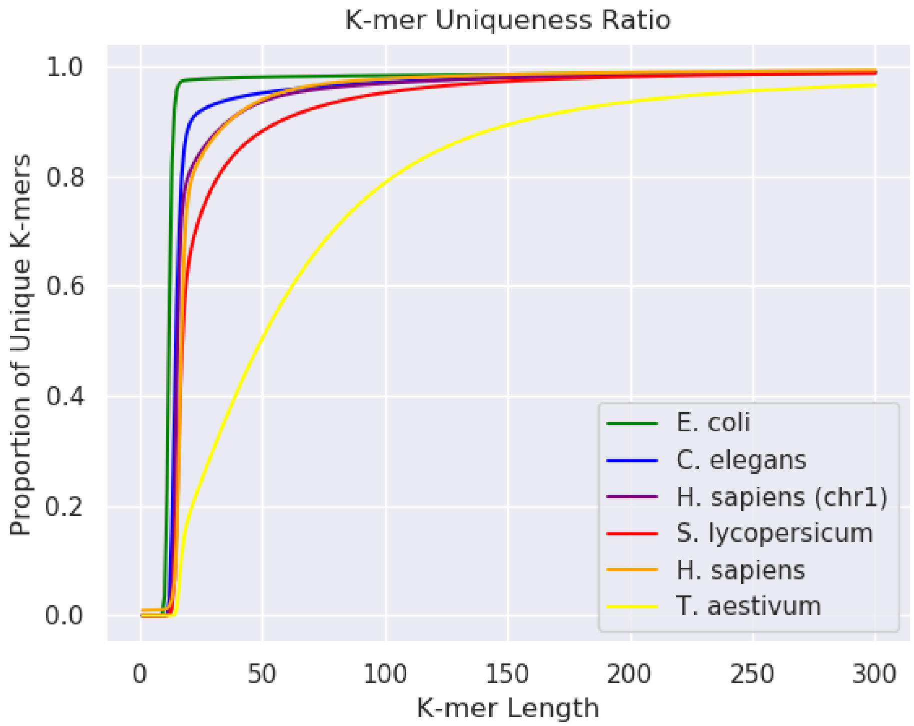

**Supplemental Figure 2. Suffix array distributions for simulated repetitive genomes.**

To illustrate the effects of repetitive sequence on the suffix array distribution, we computed the suffix array for a number of simulated sequences consisting entirely of repeats of varying length. Each sequence consists of some repeat occurring consecutively for the entirely of the sequence length (set to 100 kbp). **a)** A repeat of the sequence “CT”. **b)** A repeat of the sequence “CAT”. **c)** A repeat of the sequence “ACGT”. **d)** A repeat of the sequence “ACTTCA”. **e)** A repeat of a random length-16 sequence. **f)** A repeat of a random length-50 sequence.

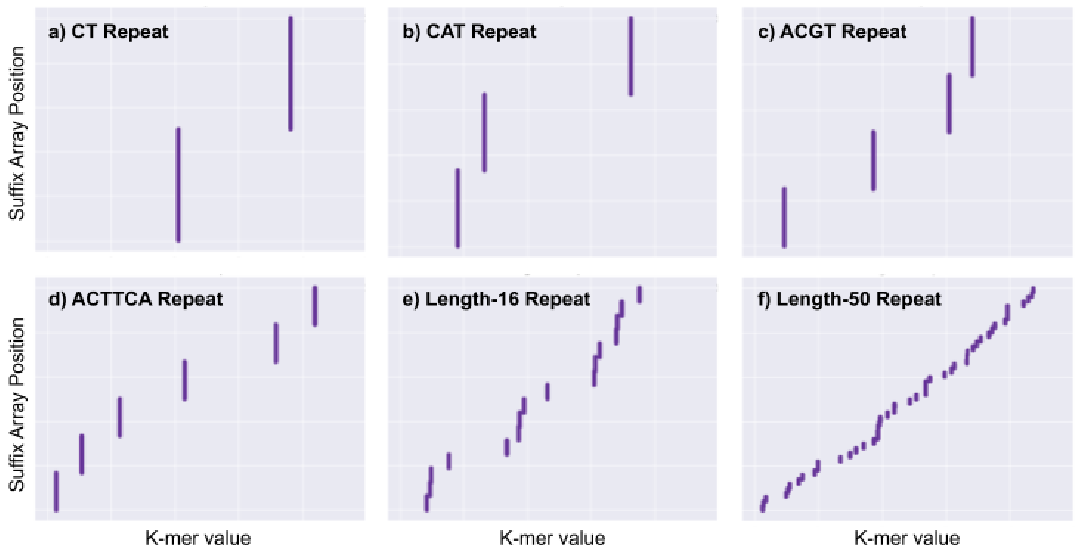

**Supplemental Figure 3. Suffix array distributions for simulated genomes with different nucleotide compositions.**

To illustrate the effects of variation in sequence content, particularly GC-content, on the suffix array distribution, we computed the suffix array for a number of simulated sequences. Each sequence consists of 100k basepairs, with each basepair independently selected from a fixed probability distribution. **a)** A sequence with 50% GC-content corresponding to an equal probability of each basepair. **b)** A sequence with 60% GC-content. **c)** A sequence with 75% GC-content. **d)** A sequence with 90% GC-content. **e)** A sequence with 100% GC-content. **f)** A sequence consisting of only C's

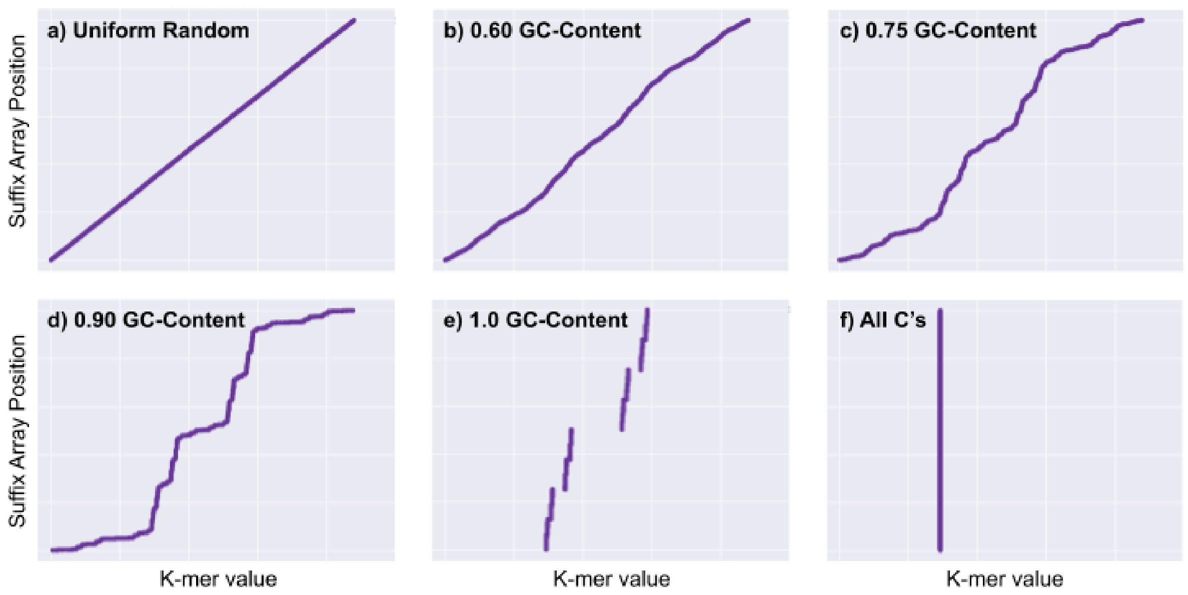


.

**Supplemental Figure 4. Distribution of the number of epochs until convergence.**

This figure shows the distribution in the number of epochs until convergence for three ANN model architectures for human chromosome 1.

a) 1024 chunks, 32Wx1L

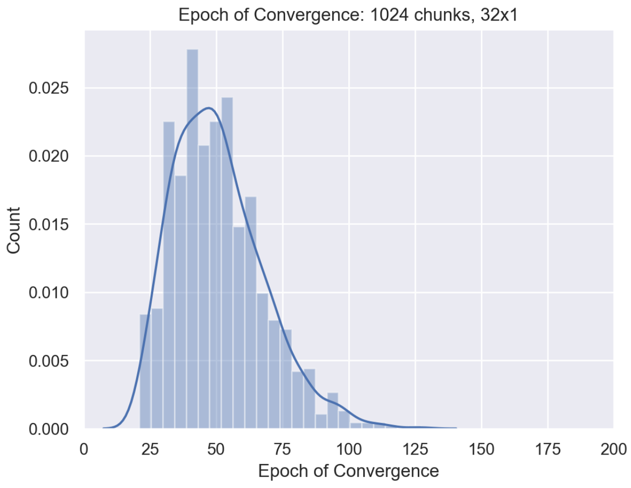


b) 16,384 chunks, 32Wx1L

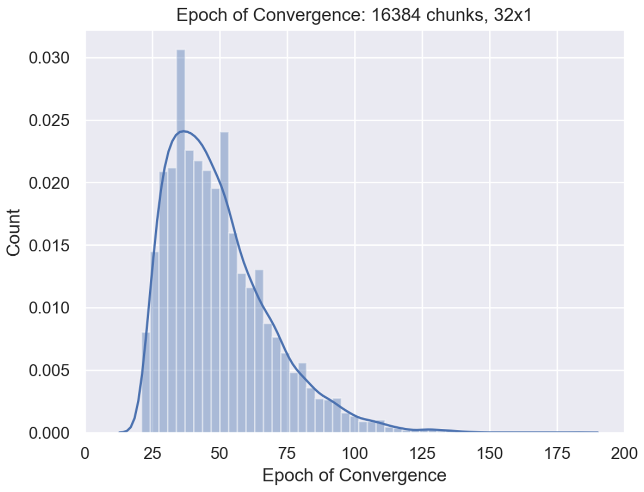


c) 16,384 chunks, 128Wx2L

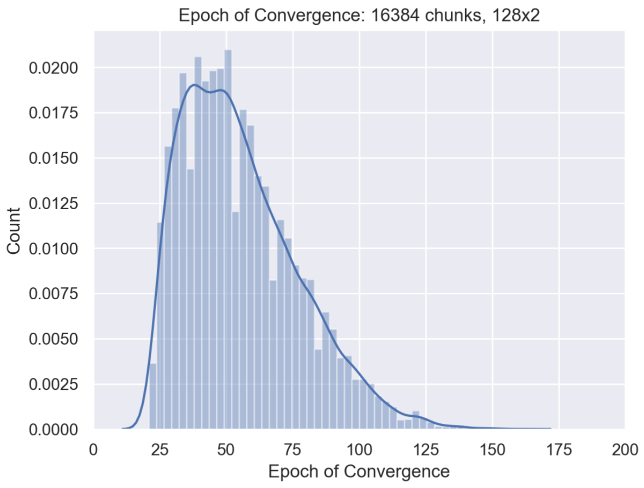

**Supplemental Figure 5. Examples of PWL models for selected bins in human chr1.**

This figure shows the functions learned by the piecewise linear data model within a few individual segments of human chromosome 1. The total number of segments in this experiment was ~16 million. Panels **a)** and **b)** show bins with the lowest mean error; Panels **c)** and **d)** show bins with average mean error; and Panels **e)** and **f)** show bins with the highest mean error. Blue shows the actual suffix array distribution and green shows the piecewise linear function for this bin.

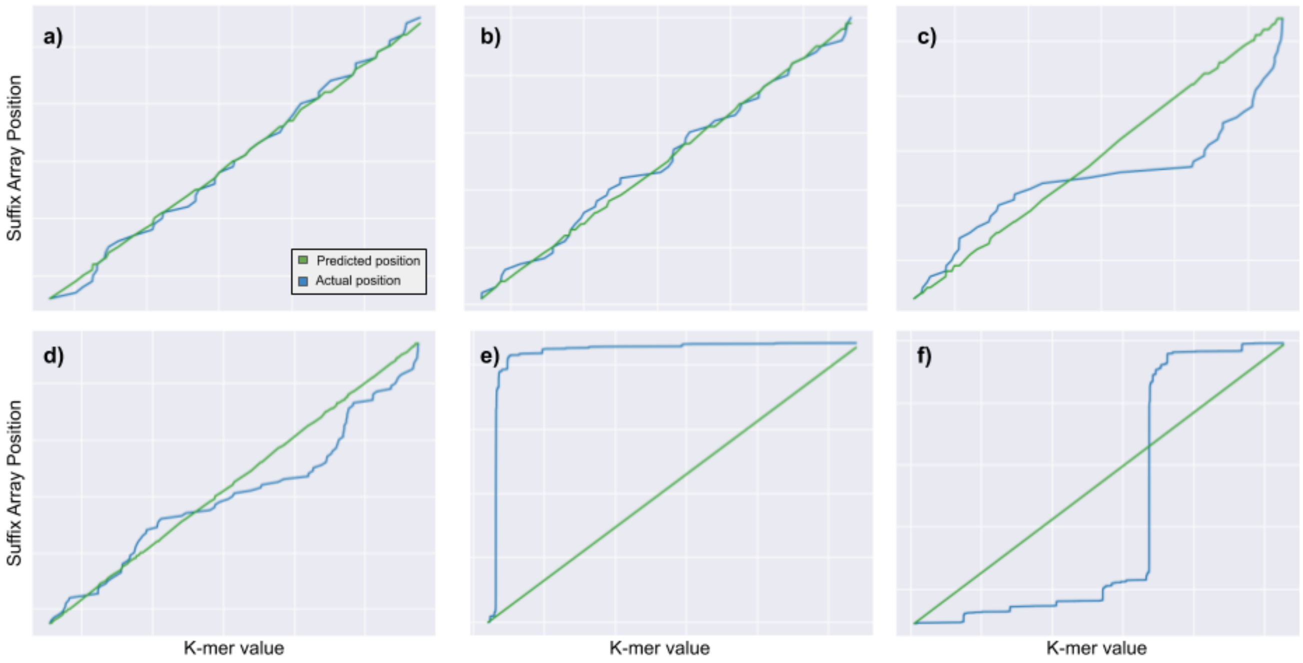

**Supplemental Figure 6. Examples of ANN performance for selected bins in human chr1.**

This figure shows some of the functions learned by the ANN models within a few bins in human chr1. The first row contains functions that are learned well (low mean error), the second row contains functions that are learned poorly (high mean error). The results shown come from the three ANN architectures highlighted in **Table 1**. Blue shows the actual suffix array distribution and green shows the ANN prediction for this bin.

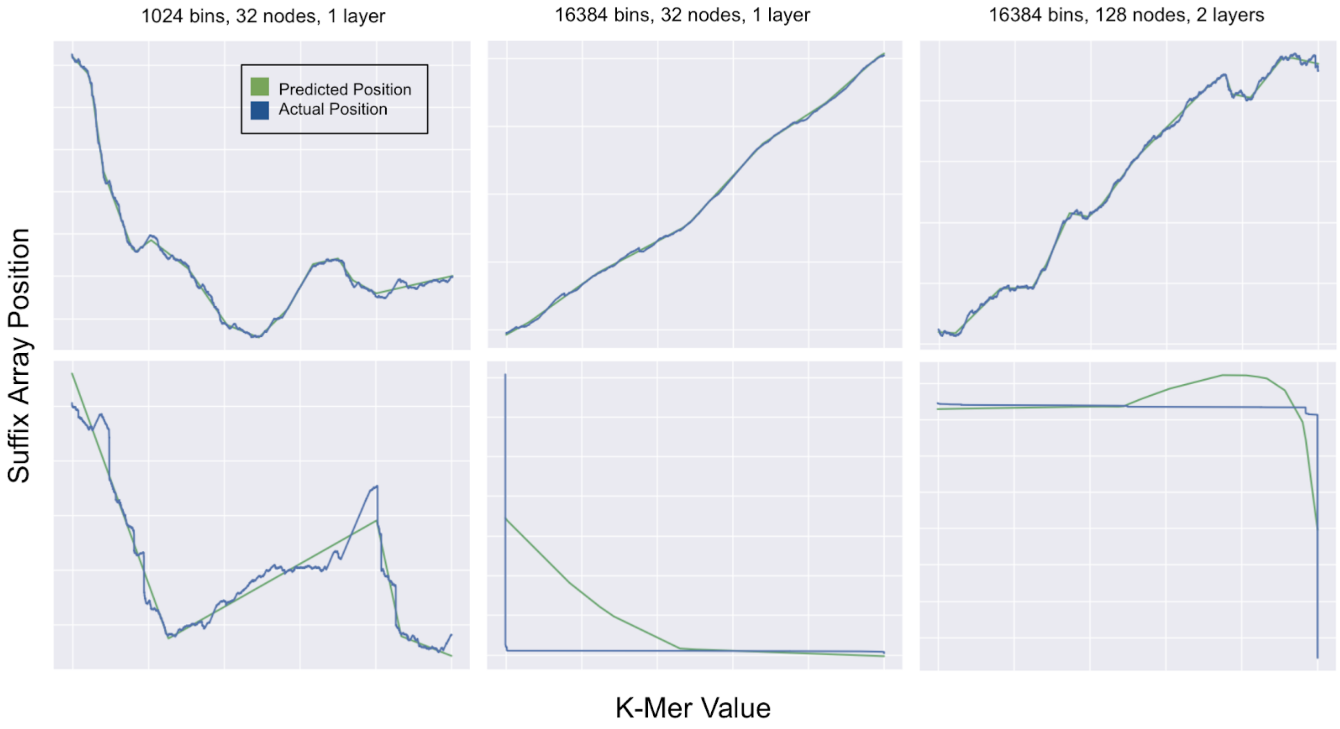

**Supplemental Note 1. Commands used for running different aligners**

**Sapling exact matching**:

```
$ suffixarray/refToSuffixArray.sh human.fa
$ src/sapling_example human.fa saFn=human.fa.sa nb=26 nq=50000000
```

**Bowtie exact matching**:

```
$ bowtiebuild human.fa human
$ bowtie -v 0 -k 1 -t human queries.fastq
```

**Mummer 4.0.0 exact matching**:

```
$ mummer -threads 1 -save human.mummer -maxmatch -l 21 human.fa \
  queries.fastq
$ time (mummer -threads 1 -load human.mummer -maxmatch -l 21 human.fa \
  queries.fastq | tee mummer.sam)
```
